# Investigating neurometabolite changes in response to median nerve stimulation

**DOI:** 10.1101/2024.05.21.593297

**Authors:** Mairi S. Houlgreave, Katherine Dyke, Adam Berrington, Stephen R. Jackson

## Abstract

**Background:** Rhythmic median nerve stimulation (MNS) at 10Hz has been shown to cause a substantial reduction in tic frequency in individuals with Tourette Syndrome. The mechanism of action is currently unknown, but is hypothesised to involve entrainment of oscillations within the sensorimotor cortex.

**Objective:** We used functional magnetic resonance spectroscopy (fMRS) to explore the dynamic effects of MNS on neurometabolite concentrations.

**Methods:** Here, we investigated the effects of rhythmic and arrhythmic 10Hz MNS on glutamate (Glu) and GABA concentrations in the contralateral sensorimotor cortex in 15 healthy controls, using a blocked fMRS design. We used a Mescher-Garwood-semi-localised by adiabatic selective refocusing (MEGA-sLASER) sequence at 7T.

**Results:** Our results show no difference in the difference-from-baseline measures between the two stimulation conditions. Looking at the effect of MNS over both conditions there is a trend for an initial increase in Glu/tCr (total creatine) followed by a decrease over time, whereas GABA/tCr decreased during each stimulation block.

**Conclusion:** These results suggest that despite entrainment of oscillations during rhythmic MNS, there are no significant differences in the tonic neuromodulatory effects of rhythmic and arrhythmic stimulation. The reduction in glutamate over the course of stimulation may reflect a decrease in glutamatergic firing due to adaptation. This may make it less likely that an involuntary movement is generated during continuous stimulation.

## 1 Introduction

Rhythmic median nerve stimulation (MNS) has been shown to result in frequency specific increases in the amplitude and phase synchronisation of neural oscillations (Houlgreave et al., 2022; Morera Maiquez et al., 2020). This modulation is restricted to the contralateral sensorimotor cortex and is not seen during arrhythmic stimulation (Houlgreave et al., 2022; Morera Maiquez et al., 2020). The mechanisms underpinning rhythmic MNS are of therapeutic interest as compared to periods of no stimulation, rhythmic application of 10Hz MNS has also been demonstrated to produce substantial reduction in tic frequency and the urge-to-tic in individuals with Tourette Syndrome (Iverson, Arbuckle, Song, et al., 2023; Iverson, Arbuckle, Ueda, et al., 2023; Maiquez et al., 2023; Morera Maiquez et al., 2020). In contrast to other non-invasive stimulation methods, specifically transcranial magnetic stimulation (TMS), MNS offers an attractive approach to modulating brain sensorimotor networks implicated in brain health conditions such as Tourette Syndrome, which could easily be adapted into a wearable therapeutic device.

Functional magnetic resonance imaging (fMRI) studies have shown that both unilateral movements and rhythmic MNS result in activation of cortical sensorimotor regions. During unilateral movements, there is activation in the contralateral sensorimotor cortex and deactivation of the ipsilateral sensorimotor cortex (Allison et al., 2000). Similarly, MNS at low frequencies (0.5-4Hz) leads to activation of the contralateral primary sensory cortex, bilateral secondary sensory cortex, and the bilateral insula (Backes et al., 2000; Ferretti et al., 2007; Manganotti et al., 2009). Furthermore, primary somatosensory cortex (S1) activation has been demonstrated to increase with stimulation frequency, although this increase plateaus at 10Hz (Ferretti et al., 2007; Kampe et al., 2000; Manganotti et al., 2009).

Chen and colleagues used an fMRI localiser task to identify a voxel located in the motor cortex which was activated by a simple hand-clenching task (Chen et al., 2017). Then, using functional magnetic resonance spectroscopy (fMRS), they demonstrated a significant increase in glutamate (Glu) and glutamine (Gln) within this voxel during the same task (Chen et al., 2017). Other fMRS studies have reported similar increases in Glu during motor tasks (Schaller et al., 2014; Volovyk & Tal, 2020). Meanwhile, a significant decrease in gamma-aminobutyric acid (GABA) was reported (Chen et al., 2017). GABA is the main inhibitory neurotransmitter in the brain, however at any timepoint, the majority of GABA in the brain forms a metabolic pool while the minority is neurotransmitter (Rae, 2014). Therefore, changes in conventional MRS-GABA collected at rest, or in a blocked fMRS designs likely reflect alterations in tonic rather than phasic inhibition (Rae, 2014; Stagg et al., 2011). On the other hand, glutamate is the main excitatory neurotransmitter in the brain, and a novel simultaneous fMRS/fMRI experiment has shown that MRS-Glu and fMRI-BOLD activation are significantly correlated over time (Ip et al., 2017). This suggests that MRS-Glu increases could reflect increases in glutamatergic neuronal firing (Ip et al., 2017), however there remains a degree of speculation regarding the origin of MRS measured signals, including for Glu (Mullins, 2018).

fMRS is a powerful approach which allows non-invasive in vivo quantification of neurometabolites. Recent studies at ultra-high field (7 T) in the visual (Boillat et al., 2020; Ip et al., 2017) and motor (Chen et al., 2017; Kolasinski et al., 2019) cortex have demonstrated that fMRS can be successfully used to detect task related alterations in the level of metabolites such as Glu and GABA. Performing ultra-high field MRS offers a higher signal-to-noise ratio (SNR) and greater spectral dispersion resulting in improved quantification. As a result, it is possible to detect contributions from peaks which overlap at lower field strengths, such as Glu and Gln, in addition to acquiring higher signal from low concentration metabolites such as GABA (Godlewska et al., 2017). The detection of GABA can be further enhanced using editing approaches which allow unambiguous assignment of GABA in the difference spectrum (Puts & Edden, 2012).

In this study we investigated the impact of repetitive MNS in healthy adults as a bridge to enhancing our knowledge into the therapeutic potential of MNS. While rhythmic MNS has been shown to influence oscillatory activity, we know little about its effects on neurometabolites, such as GABA and Glu. Given the neurometabolic changes associated with sensorimotor activation during movement (Chen et al., 2017), we hypothesise that repetitive trains of MNS at 10Hz may lead to an increase in Glu and a decrease in GABA concentration. This study aims to test this hypothesis using ultra-high field fMRS.

## 2 Methods

### 2.1 Participants

Seventeen neurologically healthy, unmedicated, adults were recruited for this study. Two were excluded prior to data collection; one due to mild claustrophobia/nausea whilst in the scanner and another due to an inability to produce a sufficient muscle twitch at a comfortable MNS intensity. The remaining sample of 15 participants completed two scanning sessions in a counterbalanced order. These sessions were spaced 10±8 days apart and the time of the session was held constant (i.e., if the first session was conducted in the morning so was the second) for all but one participant due to a change in availability. All participants were deemed right-handed using the Edinburgh Handedness inventory (Oldfield, 1971). The mean participant age was 27±5 years and 8 were female. Participant demographics can be seen in Table 1. The study received ethical approval through the University of Nottingham School of Psychology committee (Reference number: F1226, Date: 8/10/20) and all participants gave informed consent.

**Table 1.** Participant demographics for rhythmic and arrhythmic conditions.

|  | N | Sex<br>(m/f) | Age<br>(years) | MNS<br>intensity<br>(mA) | Difference in<br>intensity<br>between<br>sessions (mA) |
| --- | --- | --- | --- | --- | --- |
| Rhythmic | 15 | 7/8 | $27.5 \pm 4.8$ | $11.3 \pm 2.1$ | $0.3 \pm 2.3$ |
| Arrhythmic | 15 | 7/8 | $27.5 \pm 4.8$ | $11.4 \pm 3.0$ | |

Note – counterbalanced within subject design used. Data are presented as mean value ± sd.

### 2.2 MNS Stimulation paradigm

Stimulation was delivered to the right median nerve using a Digitimer constant current stimulator model DS7A (Digitimer Ltd, UK). The maximum compliance voltage (Vmax) was set to 400V, and the pulse width was 0.2ms. The stimulation threshold for each participant was determined to be the minimum intensity which induced a visible thumb twitch (Table 1). During the stimulation blocks, which lasted 500s, stimulation was delivered at 10Hz for 1s followed by 2s of no stimulation (MATLAB R2017a, Mathworks, Natick, MA). Stimulation was not delivered constantly for the 500s to ensure participant comfort. The three stimulation blocks were interspersed with blocks of no stimulation lasting 200s. Each participant completed one session of rhythmic stimulation and one of arrhythmic stimulation. The arrhythmic session was used to investigate whether similar neurometabolic changes occurred when stimulation had a random interpulse interval but the same average frequency (minimum interpulse interval of 0.01s) (Figure 1).

**Figure 1.**
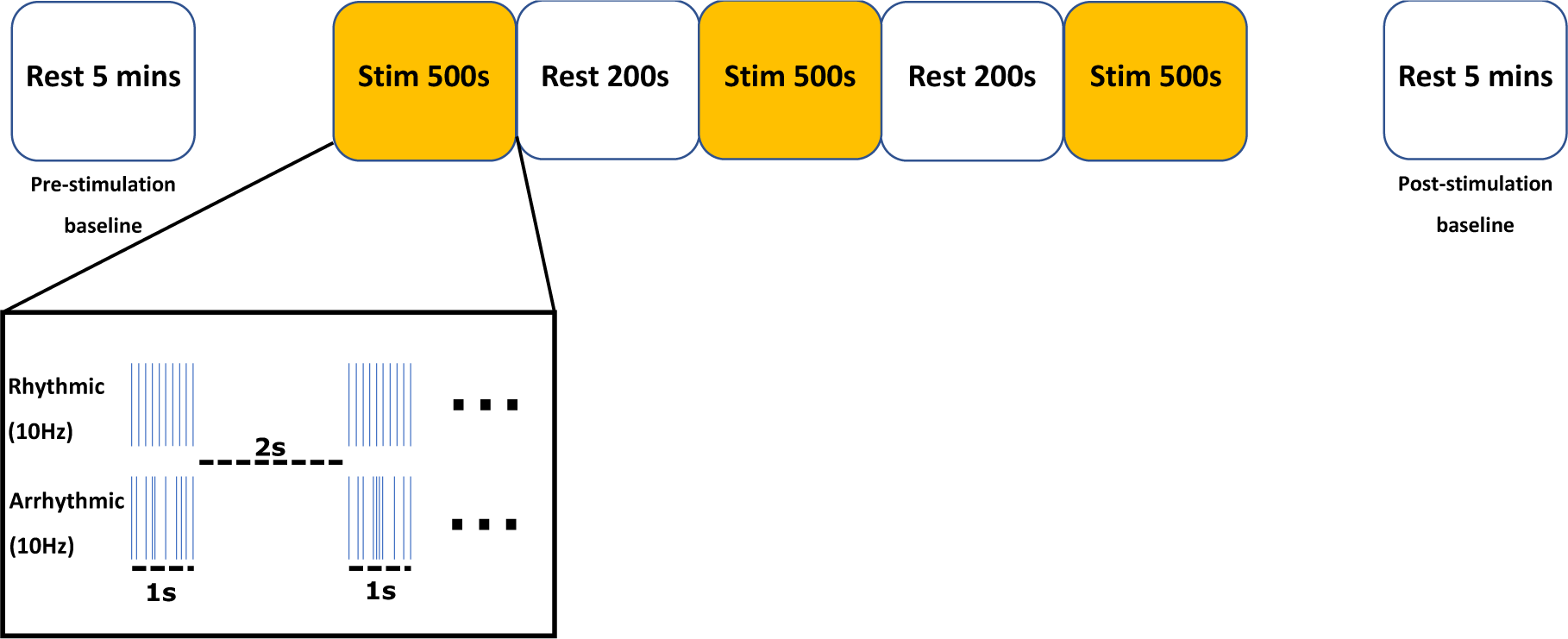
A diagram demonstrating both the trial setup and the stimulation paradigm.

### 2.3 MR acquisitions

MR data were acquired using a Philips 7 T Achieva MR scanner (Philips Healthcare, Best, The Netherlands) with a 32-channel receive array head coil. A pair of prism glasses were used to permit participants to view a nature documentary displayed on a screen outside of the scanner bore for the duration of the scan.

*T_1_*-weighted anatomical images were acquired using a MPRAGE sequence (TR/TE/TI=7.3/3.4/1000ms, FA=8°, FOV=256×256×180mm^3^, isotropic resolution=1mm^3^) for tissue segmentation (using SPM12) and planning of the MRS voxel. ^1^H MRS data were acquired from a voxel of interest (30×30×30mm^3^) placed over the contralateral hand area (Figure 2) using a MEGA-sLASER sequence optimised for GABA (TR/TE=4640/72ms, spectral width = 4kHz) (Andreychenko et al., 2012; Mescher et al., 1998). Water suppression was achieved using VAPOR (variable pulse power and optimised relaxation delay) (Tkáč et al., 1999). B^0^-shimming was performed using a vendor-provided second-order projection-based method. Further methodological details are provided in supplementary materials in accordance with recent recommendations (Lin et al., 2021).

**Figure 2.**
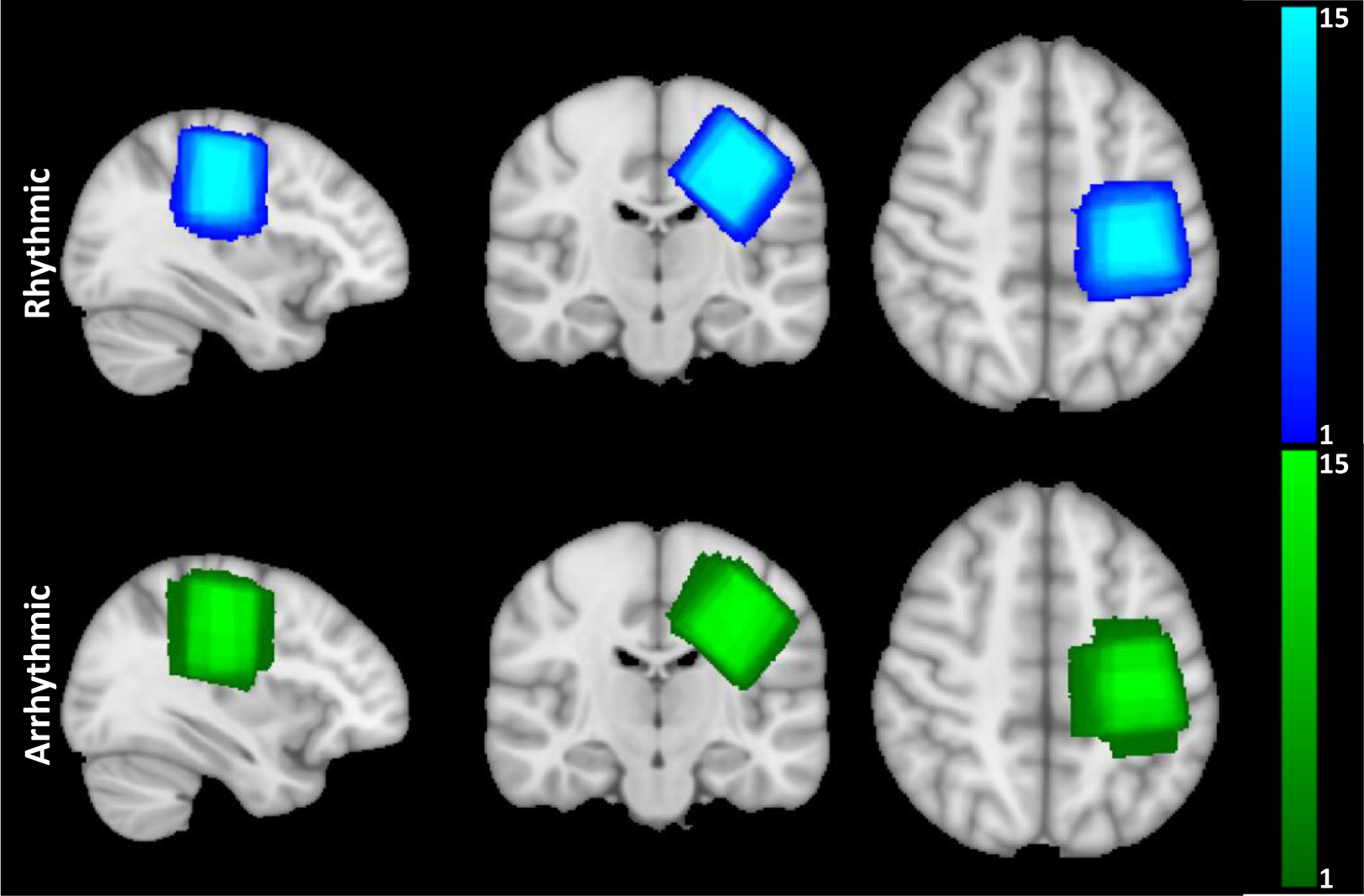
MRS voxel (30×30×30mm^3^) overlaps for the rhythmic and arrhythmic sessions centred on the hand knob of the contralateral sensorimotor cortex. Colour bars signify the number of subjects.

Three consecutive MRS scans were performed during each experimental session. The MEGA-sLASER parameters were identical for each, except for number of signals averaged and hence total scan time. Pre and post MNS scans occurred without stimulation and lasted approximately 5mins 20s and consisted of 64 transients consisting of 32 ON/OFF pairs. Scans taken during MNS in both the rhythmic and arrhythmic conditions lasted approximately 32mins 5s and consisted of a total of 410 transients consisting of 205 ON/OFF pairs. For two participants in one session, the voxel was repositioned following the pre-stimulation baseline scan due to movement at the beginning of the stimulation block.

### 2.4 Data analysis

Raw spectral data (.data/.list format) were pre-processed using an in-house MATLAB script (MATLAB R2020a, Mathworks, Natick, MA). Raw data were coil-combined and eddy-current corrected before being split into ON and OFF spectra. Spectral registration was performed to align individual transients (frequency and phase) to the mean OFF spectra for that participant (Near et al., 2015). Following alignment, individual transients were rejected if the mean square error around the Choline (Cho) peak differed from the mean by more than 3 standard deviations. The aligned ON and OFF spectra were then subtracted to create difference (DIFF) spectra.

For the fMRS acquired during stimulation, spectra were averaged over each block to create a timecourse. Blocks comprised of a spectral average of 108 transients (54 DIFF spectra) for the stimulation periods and 42 transients (21 DIFF spectra) for the rest periods. Each timecourse comprised 5 timepoints. For the pre- and post-stimulation scans we obtained 1 timepoint for each, consisting of 64 transients.

The GABA DIFF and OFF spectra were fitted in LCModel (Provencher, 2001). Spectra were fit with simulated basis spectra from 2D density matrix simulations with shaped refocusing pulse information and inter-pulse timings (Govind et al., 2015; Tkáč, 2008). The LCModel *nobase* control parameter was set to false to enable baseline fitting. The knot parameter was set to DKNTMN. The spectral range was set to 1.8-4.2ppm. The tCr (total creatine) and Glu concentrations were quantified using the LCModel concentrations in the OFF spectra, while GABA was quantified using the DIFF spectra. Concentrations are presented as a ratio relative to tCr. *A priori* exclusion criteria were if the SNR of NAA (N-acetylaspartate) was less than 40, or if the linewidth of unsuppressed water was greater than 15Hz (0.05 ppm). No participants were excluded by these criteria. The SNR was calculated using the FID-A MRS toolbox (https://github.com/CIC-methods/FID-A) (Simpson et al., 2017). One participant was excluded due to having noisy spectra (Female, 37 years, right-handed, 11.5 mA intensity for both conditions).

### 2.5 Statistical analysis

Change ratios were calculated with respect to the pre-stimulation baseline for that session (cf. Chen et al., 2017). These difference-from-baseline measures were then standardised (Z-transformed) for each participant. Difference-from-baseline measures for each block for rhythmic verses arrhythmic were compared using a paired t-test, or a Wilcoxon signed rank test where data failed the Kolmogorov-Smirnov test of normality. Difference-from-baseline measures for the combined stimulation data for each block were compared using a oneway ANOVA. The effect size was measured using Cohen’s d. To examine if difference-from-baseline values were modulated by period, a regression (GLM) analysis was conducted with stimulation (i.e., on vs. off) entered as a predictor. An additional GLM analysis was conducted with block (1-6) entered as a predictor, to investigate if there was a linear trend for Glu/tCr difference values to decrease over time.

## 3 Results

The data quality metrics of the MRS data including SNR, unsuppressed water linewidth and Cramér–Rao lower bounds (CRLBs) for Glu and GABA can be seen in Table 2. The low linewidth of the water peak implies that good shimming was achieved. Figure 3 shows the quality of both the GABA and glutamate LCModel spectra fit at the individual-level, from a representative subject, during the fMRS stimulation blocks. To check that we could reliably fit glutamine from glutamate, we ensured that the pair-wise correlation coefficient for all scans was greater than -0.5.

**Figure 3.**
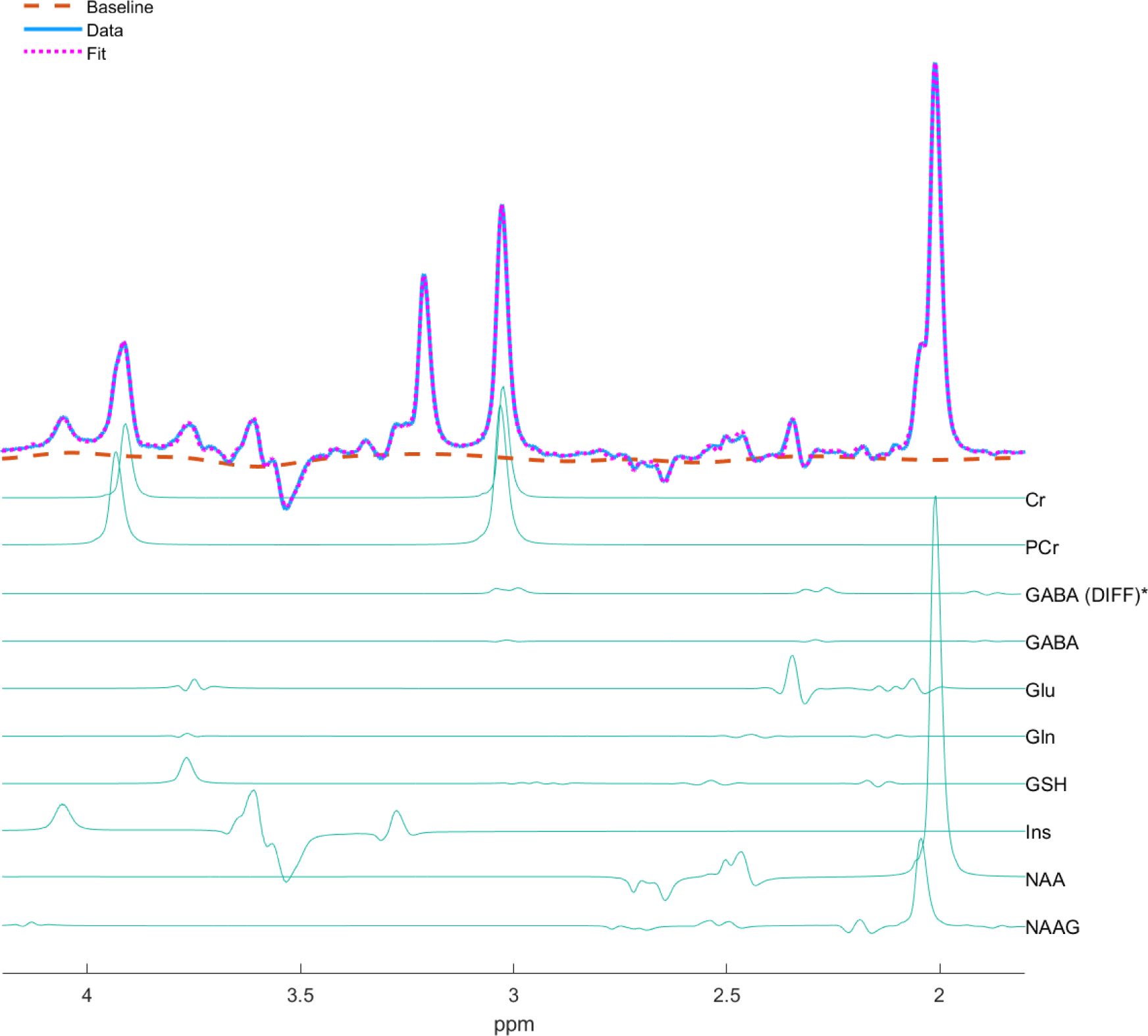
Example MEGA-sLASER OFF spectra (TE=72 ms) from an individual subject session showing the average spectra from the fMRS stimulation scan (blue) and the average LCModel fit (pink dashed line). (*GABA concentrations were measured from DIFF spectra).

**Table 2.** Data quality metrics for all scans in the rhythmic and arrhythmic conditions (N=14).

|  | Rhythmic baseline | Rhythmic post-stimulation | Rhythmic stimulation | Arrhythmic baseline | Arrhythmic post-stimulation | Arrhythmic stimulation |
| --- | --- | --- | --- | --- | --- | --- |
| Linewidth of unsuppressed water peak (Hz) | 11.1 ± 1.0 | 11.2 ± 0.9 | 10.9 ± 0.9 | 11.0 ± 1.4 | 11.3 ± 1.2 | 11.0 ± 1.0 |
| SNR of NAA | 281.8 ± 41.2 | 274.0 ± 35.4 | 693.7 ± 86.5 | 273.5 ± 51.5 | 259.4 ± 47.5 | 659.3 ± 114.0 |
| GABA CRLB (%) | 7.9 ± 2.3 | 7.9 ± 1.9 | 7.3 ± 1.3 | 7.8 ± 1.5 | 7.6 ± 1.6 | 7.6 ± 1.9 |
| Glu CRLB (%) | 3.4 ± 0.5 | 3.5 ± 0.5 | 3.4 ± 0.5 | 3.4 ± 0.6 | 3.4 ± 0.5 | 3.4 ± 0.5 |

Data are presented as mean value ± SD. SNR, signal-to-noise ratio; NAA, N-acetylaspartate; GABA, γ-aminobutyric acid; CRLB, Cramér–Rao lower bound; Glu, glutamate.

Figure 2 shows reliable positioning of the MRS voxel over the contralateral hand area for both stimulation sessions. This voxel was composed of 67 ± 3% white matter (WM), 29 ± 3% grey matter (GM) and 4 ± 2% cerebrospinal fluid (CSF). Paired samples t-tests confirmed that there were no significant differences in voxel composition for WM, GM or CSF between rhythmic and arrhythmic sessions (all p > 0.2).

Initial analyses demonstrated that the difference-from-baseline measure for rhythmic compared to arrhythmic stimulation was equivalent for both Glu/tCr (p > 0.05) or GABA/tCr (p > 0.05) ratios. Relevant means are presented in Figure 4. Therefore, rhythmic and arrhythmic data were combined to explore the effect of stimulation, regardless of pattern, on neuro-metabolite concentration.

**Figure 4.**
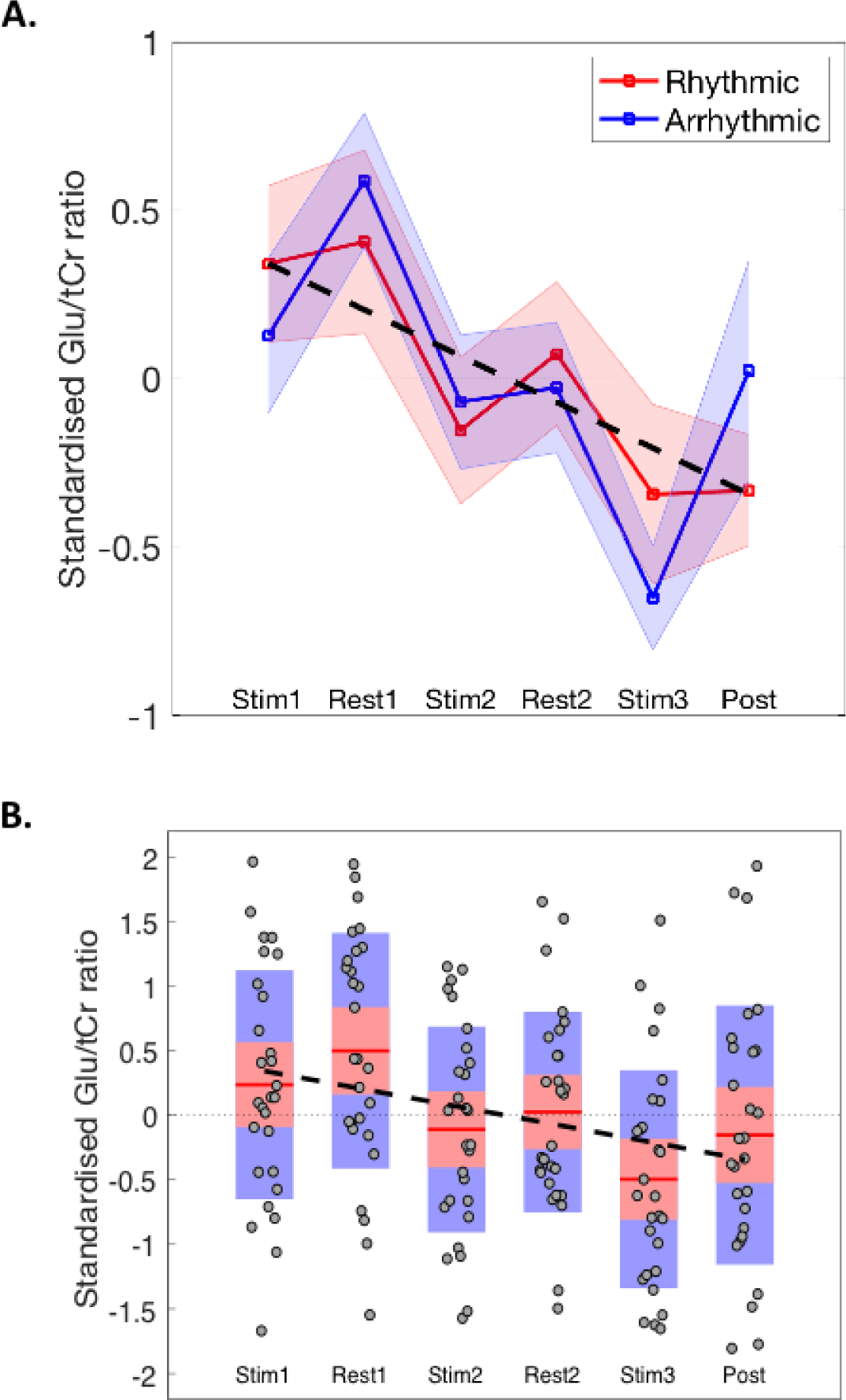
A. Difference-from-baseline measures for Glu/tCr ratio for rhythmic versus arrhythmic stimulation conditions. The shading represents the standard error of the mean (sem). The broken black line illustrates a linear line-of-best-fit for the combined data. B. Box-and-whisker plots of the combined data. Mean values are indicated by the red horizontal lines. The red and blue rectangles represent one standard deviation (red) and the 95% confidence interval (blue). Individual data points are represented by black circles. The broken black line illustrates a linear line-of-best-fit for the combined data.

### 3.1 Difference-from-baseline Glu/tCr

Inspection of Figure 4 suggests that Glu/tCr ratio difference values are modulated by stimulation (i.e., on vs. off periods) and that the magnitude of the Glu/tCr ratio difference values decrease over time. A oneway ANOVA demonstrated that there was a statistically significant effect of period (F(5,162) = 4.3, p < 0.002). To examine if Glu/tCr difference values were modulated by period, a regression (GLM) analysis was conducted with stimulation (i.e., on vs. off) entered as a predictor. This analyses demonstrated that the effect of stimulation did not reach conventional levels of statistical significance (F= 3.09, Rsq = 0.02, p = 0.08). To examine if there was a linear trend for Glu/tCr difference values to decrease over time, a second regression (GLM) analysis was conducted with block (1-6) entered as a predictor. This analyses demonstrated that the effect of block was statistically significant and confirmed that Glu/tCr decreased linearly over time (Slope = -0.14, F= 11.7, Rsq = 0.07, p < 0.001).

### 3.2 Difference-from-baseline GABA/tCr

Inspection of Figure 5 suggests that GABA/tCr ratio difference values are modulated by stimulation (i.e., on vs. off periods). A oneway ANOVA demonstrated that there was a statistically significant effect of period (F(5,162) = 5.3, p = 0.0002). To specifically examine if GABA/tCr difference values were modulated by period, a regression (GLM) analysis was conducted with stimulation (i.e., on vs. off) entered as a predictor. This analyses demonstrated that there was a strong and statistically significant effect of stimulation (F= 18.8, Rsq = 0.1, p < 0.0001). Additional analyses showed that this effect was present if Rhythmic and Arrhythmic stimulation were examined separately (minimum F = 7.0, p < 0.01).

**Figure 5.**
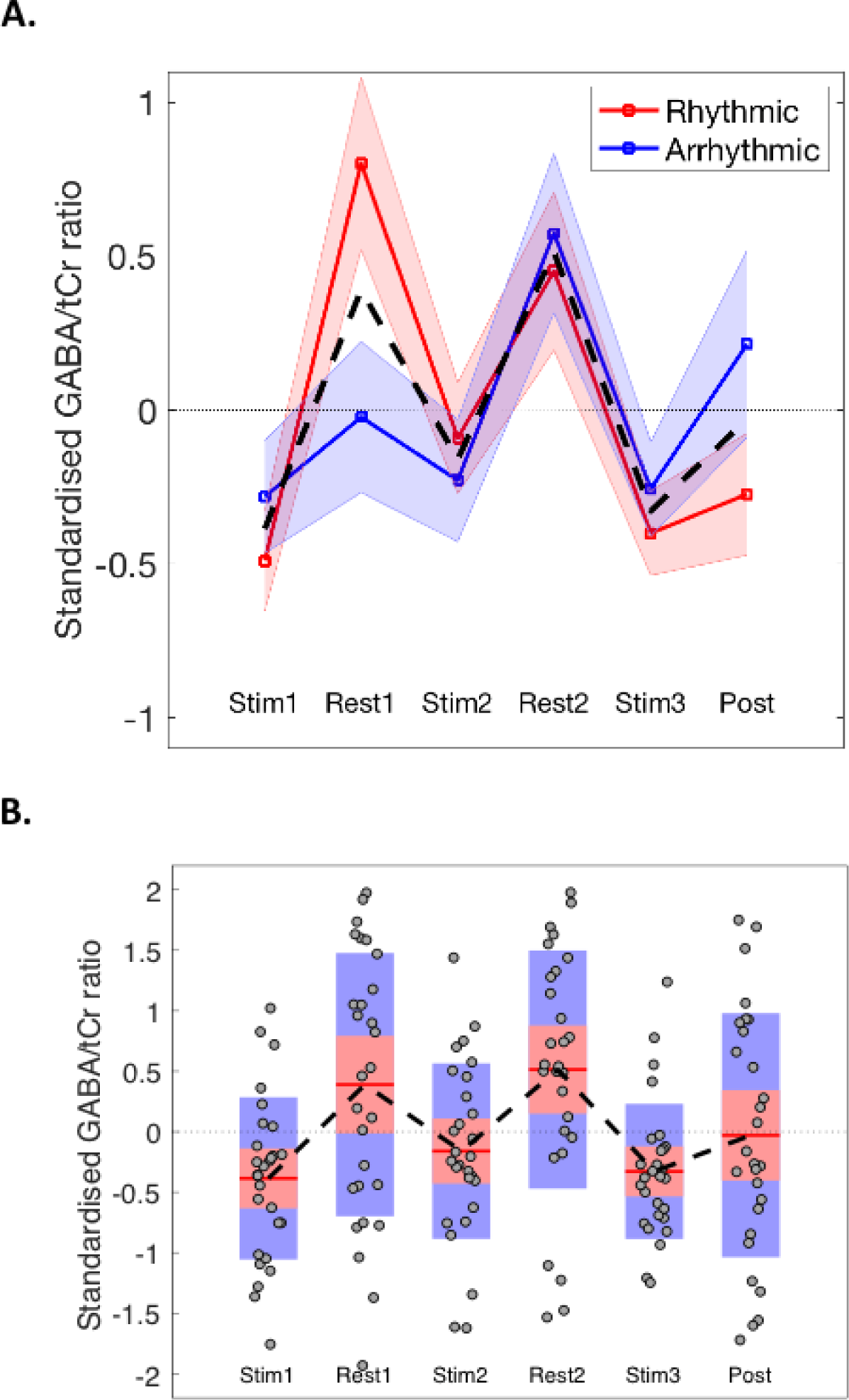
A. Difference-from-baseline measures for GABA/tCr ratio for rhythmic versus arrhythmic stimulation conditions. The shading represents the standard error of the mean (sem). The broken black line illustrates the mean values for the combined data. B. Box-and-whisker plots of the combined data. Mean values are indicated by the red horizontal lines. The red and blue rectangles represent one standard deviation (red) and the 95% confidence interval (blue). Individual data points are represented by black circles.

## 4 Discussion

This fMRS study investigated the effects of rhythmic and arrhythmic MNS on neurometabolite concentrations in the contralateral sensorimotor cortex. When overall difference-from-baseline measures were compared there were no statistically significant differences observed between the effects of rhythmic and arrhythmic stimulation on Glu or GABA concentrations. However, when we examined the effects of stimuluation irrespective of stimulation pattern (i.e., rhythmic or arrhythmic stimulation), we do see clear evidence that stimulation modulates concentrations of both Glu and GABA. Specifically, stimulation leads to a linear decrease in Glu concentration over time. By contrast, GABA concentrations decrease during stimulation but increase once again each time stimulation ceases. These effects are discussed below.

Previous studies involving a motor task have also reported increases in Glu (Chen et al., 2017; Schaller et al., 2014; Volovyk & Tal, 2020). Our finding of increased Glu during somatosensory stimulation is consistent with the findings reported by Chen et al. (2017), that Glu concentration increases during sensorimotor activity. Addtionally, in our study stimulation led to a significant reduction in GABA concentration, with higher concentrations seen during rest. These results are also in line with Chen et al. (2017), who reported a decrease in GABA during a sensorimotor hand clenching task. However, such decreases in GABA have not been reliably reported. A study by Volovyk & Tal (2020) investigated alterations in Glu and GABA during a hand clenching task (Volovyk & Tal, 2020). They reported increases in Glu during hand clenching but no change in GABA. However, it should be noted that this study was conducted at 3T rather than 7T. Similarly, a pre-clinical study demonstrated an increase in Glu but no change in GABA in the contralateral S1 of mice during electrical hind paw stimulation (Seuwen et al., 2019).

The difference between the rhythmic and arrhythmic stimulation conditions is the rhythmicity, or not, of the stimulation. Thus, both forms of stimulation contain an identical number of pulses, of the same duration and intensity, and the mean stimulation frequency is the same. We have previously demonstrated using magnetoencephalography (MEG) that rhythmic MNS increased the power and phase-synchrony of brain oscillations at the stimulation frequency within the contralateral sensorimotor area, whereas arrhythmic stimulation did not (Houlgreave et al., 2022). Furthermore, in a recent, as yet unpublished, study we found that both rhythmic and arrhythmic MNS delivered to the right wrist leads to a significant increase in fMRI BOLD response in contralateral primary somatosensory cortex, bilateral secondary somatosensory cortex, and bilateral insula cortex. For both patterns of stimulation, each single MNS pulse will lead to synchronous firing within a population of neurons within the contralateral somatosensory cortex; meaning excited pyramidal cells will be releasing glutamate phasically in synchrony. Importantly, Glu concentrations have been shown to be lower during repetitive compared to novel trials (Apšvalka et al., 2015), and in the currently study we found that Glu levels show an initial increase but then reduce linearly over successive blocks. Therefore, it may be that in both conditions Glu reduces over time due to the repetitive nature of the stimulation. This adaptation has not been shown in previous MRS studies involving motor tasks (Chen et al., 2017; Schaller et al., 2014; Volovyk & Tal, 2020), but here the stimulation is at a high frequency and externally driven.

Our hypotheses for this study did not consider entrainment effects of the rhythmic stimulation on sensorimotor oscillations (Houlgreave et al., 2022; Morera Maiquez et al., 2020). GABA is thought to be involved in the generation of synchronised oscillations (Gonzalez-Burgos & Lewis, 2008). Previous electrophysiological studies have demonstrated a clear relationship between GABA and oscillations within the sensorimotor cortex. Elevation of the effects of extracellular GABA, using transporter blocker tiagabine and extra-synaptic positive allosteric modulator gaboxadol, resulted in an increase in the power of all frequency bands up to and including beta oscillations (Nutt et al., 2015). Higher levels of resting MRS-GABA in the motor cortex have been associated with higher power during the post-movement beta rebound (Gaetz et al., 2011). Given this relationship between GABA and oscillatory activity, an increase in GABA related to entrainment may have been expected in the rhythmic condition (Spooner et al., 2022). However, the effects of entrainment are, by definition, restricted to the frequency of stimulation. During rhythmic MNS desynchronisation of frequencies within the 8-30Hz range was evident (Houlgreave et al., 2022; Morera Maiquez et al., 2020). Moreover, any inhibitory effects of entrainment through MNS will be phasic. Phasic inhibition relates to synaptic GABA release leading to short-lived neuronal hyperpolarisation (Brickley & Mody, 2012). By contrast, extra-synaptic GABA binding leads to a more long-lived tonic inhibition (Brickley & Mody, 2012). There is evidence to suggest that there is no association between MRS-GABA and TMS measures which are thought to reflect activity involving synaptic GABA in adults (Dyke et al., 2017; Stagg et al., 2011; Tremblay et al., 2013). This suggests that MRS-GABA may be a measure of tonic rather than phasic inhibition. As such, increases in phasic GABAergic activity relating to MNS induced entrainment are unlikely to be quantifiable through MRS. However, it is worth noting that there is evidence for an association between MRS-GABA and TMS measures in paediatric populations (Harris et al., 2021). In the current study we clearly demonstrate that GABA levels are substantially decreased during stimulation but are elevated once again during periods of no stimulation. Furthermore, this repeated reduction of GABA levels during stimulation is observed for *both* rhythmic and arrhythmic stimulation indication that any modulation of GABA levels is not frequency specific and is unlikely to be linked to the phasic release of GABA in the form of neurotransmitter.

One limitation of this study is that the stimulation was not constant during the stimulation blocks. Instead, the stimulation was delivered for 1 second followed by 2 seconds of no stimulation. This choice was made to ensure participant comfort during the experiment. However, data from previous studies shows a gradual return of GABA concentration to baseline following a movement task (Chen et al., 2017), and a gradual increase in GABA concentration during the task even when there were brief pauses between movements (Kolasinski et al., 2019). Another limitation is that the size and therefore SNR of the pre-stimulation baseline, rest blocks, stimulation blocks and post-stimulation blocks were not equivalent. However, the Cramér–Rao lower bounds for Glu and GABA fitting were not significantly different across all scans. Recent dynamic fitting approaches for MRS, incorporating temporal modelling of the stimulation periods, may offer improved ability to detect changes compared with block-averaging (Clarke et al., 2024; Tal, 2023).

## 5 Conclusions

To conclude, this research demonstrates that there is a trend for an initial rise in glutamate concentration followed by a decrease over repeated trials, and decreases in GABA during stimulation which recovered during rest, regardless of the pattern of stimulation. Furthermore, the neuromodulatory effects of rhythmic and arrhythmic stimulation were similar despite the entrainment seen previously with rhythmic stimulation (Houlgreave et al., 2022). Recent evidence also suggests that both rhythmic and arrhythmic stimulation may be effective in reducing tics in Tourette Syndrome (Iverson, Arbuckle, Song, et al., 2023; Iverson, Arbuckle, Ueda, et al., 2023; Maiquez et al., 2023; Morera Maiquez et al., 2020). We suggested previously that if both rhythmic and arrhythmic stimulation were effective in reducing tic frequency in Tourette Syndrome, then the beneficial effects might be due to a sustained decrease in sensorimotor noise after stimulation, due to the synchronous firing of activated neuronal populations associated with each pulse of stimulation (Houlgreave et al., 2022). This might be accompanied by alterations in tonic levels of neurometabolite concentrations. Here we show a reduction in glutamate over time, which may reflect a decrease in glutamatergic firing due to adaptation to the continuous stimulation (Apšvalka et al., 2015; Ip et al., 2017). This may make it less likely that an involuntary movement is generated during continuous stimulation. As an offline reduction in tic severity has been reported following 4-weeks of rhythmic MNS (Maiquez et al., 2023), it would be interesting to explore the offline changes in neurometabolites following prolonged periods of stimulation.

## Data and Code Availability

Magnetic resonance spectroscopy data can be made available on request if a formal data sharing agreement is in place. The MATLAB code for MNS delivery is available on OSF https://osf.io/c6pwa/

## Author Contributions

**MH.** Conceptualisation, Methodology, Formal Analysis, Investigation, Writing – Original Draft

**KD.** Conceptualisation, Methodology, Investigation, Writing – Original Draft, Writing – Review & Editing

**AB.** Formal Analysis, Investigation, Writing – Review & Editing

**SJ.** Conceptualisation, Methodology, Formal Analysis, Writing – Review & Editing, Supervision, Funding Acquisition

## Abbreviations

Cho: choline
CRLB: Cramér–Rao lower bound
CSF: cerebrospinal fluid
DIFF: difference spectra
fMRI: functional magnetic resonance imaging
fMRS: functional magnetic resonance spectroscopy
GABA: gamma-aminobutyric acid
GLM: general linear model
Gln: glutamine
Glu: glutamate
GM: grey matter
MEG: magnetoencephalography
MEGA-sLASER: Mescher-Garwood-semi-localised by adiabatic selective refocusing
MNS: median nerve stimulation
NAA: N-acetylaspartate
S1: primary somatosensory cortex
sem: standard error of the mean
SNR: signal to noise ratio
tCr: total creatine
TE: echo time
TI: inversion time
TMS: transcranial magnetic stimulation
TR: Repetition time
VAPOR: variable pulse power and optimised relaxation delay
Vmax: maximum compliance voltage
WM: white matter

## Acknowledgements

Data collection for this paper was supported by research grants from NIHR Nottingham Biomedical Research Centre, Medical Research Council (MRC) (T032588) and Tourette Association of America (TAA). The views expressed are those of the authors and not necessarily those of the NHS, the NIHR, MRC, TAA or the Department of Health.

## References

Allison, J. D., Meador, K. J., Loring, D. W., Figueroa, R. E., & Wright, J. C. (2000). Functional MRI cerebral activation and deactivation during finger movement. Neurology, 54(1), 135–135. 10.1212/WNL.54.1.135

Andreychenko, A., Boer, V. O., Arteaga De Castro, C. S., Luijten, P. R., & Klomp, D. W. J. (2012). Efficient spectral editing at 7 T: GABA detection with MEGA-sLASER. Magnetic Resonance in Medicine, 68(4), 1018–1025. 10.1002/MRM.24131

Apšvalka, D., Gadie, A., Clemence, M., & Mullins, P. G. (2015). Event-related dynamics of glutamate and BOLD effects measured using functional magnetic resonance spectroscopy (fMRS) at 3T in a repetition suppression paradigm. NeuroImage, 118, 292–300. 10.1016/j.neuroimage.2015.06.015

Backes, W. H., Mess, W. H., Van Kranen-Mastenbroek, V., & Reulen, J. P. H. (2000). Somatosensory cortex responses to median nerve stimulation: fMRI effects of current amplitude and selective attention. Clinical Neurophysiology, 111(10), 1738–1744. 10.1016/S1388-2457(00)00420-X

Boillat, Y., Xin, L., van der Zwaag, W., & Gruetter, R. (2020). Metabolite concentration changes associated with positive and negative BOLD responses in the human visual cortex: A functional MRS study at 7 Tesla. Journal of Cerebral Blood Flow and Metabolism, 40(3), 488–500. 10.1177/0271678X19831022

Brickley, S. G., & Mody, I. (2012). Extrasynaptic GABA A Receptors: Their Function in the CNS and Implications for Disease. In Neuron (Vol. 73, Issue 1, pp. 23– 34). 10.1016/j.neuron.2011.12.012

Chen, C., Sigurdsson, H. P., Pépés, S. E., Auer, D. P., Morris, P. G., Morgan, P. S., Gowland, P. A., & Jackson, S. R. (2017). Activation induced changes in GABA: Functional MRS at 7 T with MEGA-sLASER. NeuroImage, 156, 207–213. 10.1016/j.neuroimage.2017.05.044

Clarke, W. T., Ligneul, C., Cottaar, M., Ip, I. B., & Jbabdi, S. (2024). Universal dynamic fitting of magnetic resonance spectroscopy. Magnetic Resonance in Medicine. 10.1002/MRM.30001

Dyke, K., Pépés, S. E., Chen, C., Kim, S., Sigurdsson, H. P., Draper, A., Husain, M., Nachev, P., Gowland, P. A., Morris, P. G., & Jackson, S. R. (2017). Comparing GABA-dependent physiological measures of inhibition with proton magnetic resonance spectroscopy measurement of GABA using ultra-high-field MRI. NeuroImage, 152, 360–370. 10.1016/j.neuroimage.2017.03.011

Ferretti, A., Babiloni, C., Arienzo, D., Del Gratta, C., Rossini, P. M., Tartaro, A., & Romani, G. L. (2007). Cortical brain responses during passive nonpainful median nerve stimulation at low frequencies (0.5-4 Hz): An fMRI study. Human Brain Mapping, 28(7), 645–653. 10.1002/hbm.20292

Gaetz, W., Edgar, J. C., Wang, D. J., & Roberts, T. P. L. (2011). Relating MEG measured motor cortical oscillations to resting γ-Aminobutyric acid (GABA) concentration. NeuroImage, 55(2), 616–621. 10.1016/j.neuroimage.2010.12.077

Godlewska, B. R., Clare, S., Cowen, P. J., & Emir, U. E. (2017). Ultra-high-field magnetic resonance spectroscopy in psychiatry. Frontiers in Psychiatry, 8(JUL), 260410. 10.3389/FPSYT.2017.00123/BIBTEX

Gonzalez-Burgos, G., & Lewis, D. A. (2008). GABA neurons and the mechanisms of network oscillations: Implications for understanding cortical dysfunction in schizophrenia. In Schizophrenia Bulletin (Vol. 34, Issue 5, pp. 944–961). 10.1093/schbul/sbn070

Govind, V., Young, K., & Maudsley, A. A. (2015). Corrigendum: proton NMR chemical shifts and coupling constants for brain metabolites. Govindaraju V, Young K, Maudsley AA, NMR Biomed. 2000; 13: 129–153. NMR in Biomedicine, 28(7), 923–924. 10.1002/NBM.3336

Harris, A. D., Gilbert, D. L., Horn, P. S., Crocetti, D., Cecil, K. M., Edden, R. A. E., Huddleston, D. A., Mostofsky, S. H., & Puts, N. A. J. (2021). Relationship between GABA levels and task-dependent cortical excitability in children with attention-deficit/hyperactivity disorder. Clinical Neurophysiology, 132(5), 1163–1172. 10.1016/j.clinph.2021.01.023

Houlgreave, M. S., Morera Maiquez, B., Brookes, M. J., & Jackson, S. R. (2022). The oscillatory effects of rhythmic median nerve stimulation. NeuroImage, 251. 10.1016/J.NEUROIMAGE.2022.118990

Ip, I. B., Berrington, A., Hess, A. T., Parker, A. J., Emir, U. E., & Bridge, H. (2017). Combined fMRI-MRS acquires simultaneous glutamate and BOLD-fMRI signals in the human brain. NeuroImage, 155(December 2016), 113–119. 10.1016/j.neuroimage.2017.04.030

Iverson, A. M., Arbuckle, A. L., Song, D. Y., Bihun, E. C., & Black, K. J. (2023). Median Nerve Stimulation for Treatment of Tics: A 4-Week Open Trial with Ecological Momentary Assessment. Journal of Clinical Medicine, 12(7). 10.3390/jcm12072545

Iverson, A. M., Arbuckle, A. L., Ueda, K., Song, D. Y., Bihun, E. C., Koller, J. M., Wallendorf, M., & Black, K. J. (2023). Median Nerve Stimulation for Treatment of Tics: Randomized, Controlled, Crossover Trial. Journal of Clinical Medicine, 12(7), 2514. 10.3390/JCM12072514/S1

Kampe, K. K. W., Jones, R. A., & Auer, D. P. (2000). Frequency dependence of the functional MRI response after electrical median nerve stimulation. Human Brain Mapping, 9(2), 106–114. 10.1002/(SICI)1097-0193(200002)9:2<106::AID-HBM5>3.0.CO;2-Y

Kolasinski, J., Hinson, E. L., Divanbeighi Zand, A. P., Rizov, A., Emir, U. E., & Stagg, C. J. (2019). The dynamics of cortical GABA in human motor learning. The Journal of Physiology, 597(1), 271–282. 10.1113/JP276626

Lin, A., Andronesi, O., Bogner, W., Choi, I. Y., Coello, E., Cudalbu, C., Juchem, C., Kemp, G. J., Kreis, R., Krššák, M., Lee, P., Maudsley, A. A., Meyerspeer, M., Mlynarik, V., Near, J., Öz, G., Peek, A. L., Puts, N. A., Ratai, E. M., … Mullins, P. G. (2021). Minimum Reporting Standards for in vivo Magnetic Resonance Spectroscopy (MRSinMRS): Experts’ consensus recommendations. NMR in Biomedicine, 34(5), e4484. 10.1002/NBM.4484

Maiquez, B. M., Smith, C., Dyke, K., Chou, C. P., Kasbia, B., McCready, C., Wright, H., Jackson, J. K., Farr, I., Badinger, E., Jackson, G. M., & Jackson, S. R. (2023). A double-blind, sham-controlled, trial of home-administered rhythmic 10-Hz median nerve stimulation for the reduction of tics, and suppression of the urge-to-tic, in individuals with Tourette syndrome and chronic tic disorder. Journal of Neuropsychology, 17(3), 540–563. 10.1111/JNP.12313

Manganotti, P., Formaggio, E., Storti, S. F., Avesani, M., Acler, M., Sala, F., Magon, S., Zoccatelli, G., Pizzini, F., Alessandrini, F., Fiaschi, A., & Beltramello, A. (2009). Steady-state activation in somatosensory cortex after changes in stimulus rate during median nerve stimulation. Magnetic Resonance Imaging, 27(9), 1175–1186. 10.1016/j.mri.2009.05.009

Mescher, M., Merkle, H., Kirsch, J., Garwood, M., & Gruetter, R. (1998). Simultaneous in vivo spectral editing and water suppression. 10.1002/(SICI)1099-1492(199810)11:6

Morera Maiquez, B., Sigurdsson, H. P., Dyke, K., Clarke, E., McGrath, P., Pasche, M., Rajendran, A., Jackson, G. M., & Jackson, S. R. (2020). Entraining Movement-Related Brain Oscillations to Suppress Tics in Tourette Syndrome. Current Biology, 30(12), 2334–2342.e3. 10.1016/j.cub.2020.04.044

Mullins, P. G. (2018). Towards a theory of functional magnetic resonance spectroscopy (fMRS): A meta-analysis and discussion of using MRS to measure changes in neurotransmitters in real time. Scandinavian Journal of Psychology, 59(1), 91–103. 10.1111/SJOP.12411

Near, J., Edden, R., Evans, C. J., Paquin, R., Harris, A., & Jezzard, P. (2015). Frequency and phase drift correction of magnetic resonance spectroscopy data by spectral registration in the time domain. Magnetic Resonance in Medicine, 73(1), 44–50. 10.1002/mrm.25094

Nutt, D., Wilson, S., Lingford-Hughes, A., Myers, J., Papadopoulos, A., & Muthukumaraswamy, S. (2015). Differences between magnetoencephalographic (MEG) spectral profiles of drugs acting on GABA at synaptic and extrasynaptic sites: A study in healthy volunteers. Neuropharmacology, 88, 155–163. 10.1016/j.neuropharm.2014.08.017

Oldfield, R. C. (1971). The assessment and analysis of handedness: The Edinburgh inventory. Neuropsychologia, 9(1), 97–113. 10.1016/0028-3932(71)90067-4

Provencher, S. W. (2001). Automatic quantitation of localized in vivo 1H spectra with LCModel. NMR in Biomedicine, 14(4), 260–264. 10.1002/nbm.698

Puts, N. A. J., & Edden, R. A. E. (2012). In vivo magnetic resonance spectroscopy of GABA: A methodological review. In Progress in Nuclear Magnetic Resonance Spectroscopy (Vol. 60, pp. 29–41). 10.1016/j.pnmrs.2011.06.001

Rae, C. D. (2014). A guide to the metabolic pathways and function of metabolites observed in human brain 1H magnetic resonance spectra. Neurochemical Research, 39(1), 1–36. 10.1007/S11064-013-1199-5/FIGURES/9

Schaller, B., Xin, L., O’Brien, K., Magill, A. W., & Gruetter, R. (2014). Are glutamate and lactate increases ubiquitous to physiological activation? A 1H functional MR spectroscopy study during motor activation in human brain at 7Tesla. NeuroImage, 93(P1), 138–145. 10.1016/j.neuroimage.2014.02.016

Seuwen, A., Schroeter, A., Grandjean, J., Schlegel, F., & Rudin, M. (2019). Functional spectroscopic imaging reveals specificity of glutamate response in mouse brain to peripheral sensory stimulation. Scientific Reports 2019 9:1, 9(1), 1–9. 10.1038/s41598-019-46477-1

Simpson, R., Devenyi, G. A., Jezzard, P., Hennessy, T. J., & Near, J. (2017). Advanced processing and simulation of MRS data using the FID appliance (FID-A)—An open source, MATLAB-based toolkit. Magnetic Resonance in Medicine, 77(1), 23–33. 10.1002/mrm.26091

Spooner, R. K., Wiesman, A. I., & Wilson, T. W. (2022). Peripheral Somatosensory Entrainment Modulates the Cross-Frequency Coupling of Movement-Related Theta-Gamma Oscillations. https://Home.Liebertpub.Com/Brain, 12(6), 524–537. 10.1089/BRAIN.2021.0003

Stagg, C. J., Bestmann, S., Constantinescu, A. O., Moreno Moreno, L., Allman, C., Mekle, R., Woolrich, M., Near, J., Johansen-Berg, H., & Rothwell, J. C. (2011). Relationship between physiological measures of excitability and levels of glutamate and GABA in the human motor cortex. Journal of Physiology, 589(23), 5845–5855. 10.1113/jphysiol.2011.216978

Tal, A. (2023). The future is 2D: spectral-temporal fitting of dynamic MRS data provides exponential gains in precision over conventional approaches. Magnetic Resonance in Medicine, 89(2), 499–507. 10.1002/MRM.29456

Tkáč, I. (2008). Refinement of simulated basis set for LCModel analysis [abstr]. Proceedings of the Sixteenth Meeting of the International Society for Magnetic Resonance in Medicine.

Tkáč, I., Starčuk, Z., Choi, I. Y., & Gruetter, R. (1999). In vivo 1H NMR spectroscopy of rat brain at 1 ms echo time. Magnetic Resonance in Medicine, 41(4), 649–656. 10.1002/(SICI)1522-2594(199904)41:4<649::AID-MRM2>3.0.CO;2-G

Tremblay, S., Beaulé, V., Proulx, S., de Beaumont, L., Marjá nska, M., Doyon, J., Pascual-Leone, A., Lassonde, M., Théoret, H., Beaumont, de L., & Rela, T. H. (2013). Relationship between transcranial magnetic stimulation measures of intracortical inhibition and spectroscopy measures of GABA and glutamate+glutamine. J Neurophysiol, 109, 1343–1349. 10.1152/jn.00704.2012.-Trans

Volovyk, O., & Tal, A. (2020). Increased Glutamate concentrations during prolonged motor activation as measured using functional Magnetic Resonance Spectroscopy at 3T. NeuroImage, 223(September). 10.1016/j.neuroimage.2020.117338

