## Supplementary material for "Investigating neurometabolite changes in response to median nerve stimulation"

**Supplementary Material**: MRSinMRS checklist

| 1. Hardware | |
|---|---|
| a. Field strength [T] | 7 T |
| b. Manufacturer | Philips Achieva |
| c. Model (software version if available) | Achieva (R5.1.7) |
| d. RF coils: nuclei (transmit/receive), number of channels, type, body part | 32 channel head coil |
| e. Additional hardware | N/A |
| **2. Acquisition** | |
| a. Pulse sequence | MEGA-sLASER |
| b. Volume of Interest (VOI) locations | Sensorimotor cortex |
| c. Nominal VOI size [$cm^3$, $mm^3$] | 3 x 3 x 3 $cm^3$ |
| d. Repetition Time (TR), Echo Time (TE) [ms] | TR 4640 ms, TE=72ms |
| e. Total number of Excitations or acquisitions per spectrum<br><br>In time series for kinetic studies<br><br>i. Number of Averaged spectra (NA) per time-point<br>ii. Averaging method (e.g. block-wise or moving average)<br>iii. Total number of spectra (acquired / in time-series) | Block-wise averaging.<br><br>PRE and POST: 64 excitations<br><br>STIM BLOCK: 108 excitations<br><br>REST BLOCK: 42 excitations |
| f. Additional sequence parameters (spectral width in Hz, number of spectral points, frequency offsets)<br><br>If STEAM:, Mixing Time (TM)<br><br>If MRSI: 2D or 3D, FOV in all directions, matrix size, acceleration factors, sampling method | SW = 4 kHz<br><br>NP=? |
| g. Water Suppression Method | VAPOR |
| h. Shimming Method, reference peak, and thresholds for "acceptance of shim" chosen | Vendor-provided projection-based shimming to second order |
| i. Triggering or motion correction method<br><br>(respiratory, peripheral, cardiac triggering, incl. device used and delays) | N/A |
| **3. Data analysis methods and outputs** | |

| | |
|---|---|
| a. Analysis software | LCModel |
| b. Processing steps deviating from quoted reference or product | Spectral registration to align to mean OFF spectrum.<br><br>Rejection of transients with MSE (mean squared error) from Cho peak. |
| c. Output measure<br><br>(e.g. absolute concentration, institutional units, ratio) | Ratio to total creatine |
| d. Quantification references and assumptions, fitting model assumptions | No relaxation corrections performed. |
| **4. Data Quality** | |
| a. Reported variables<br><br>(SNR, Linewidth (with reference peaks)) | *SNR and linewidth provided in Table 2* |
| b. Data exclusion criteria | *SNR < 40 and water linewidth > 15Hz* |
| c. Quality measures of postprocessing Model fitting (e.g. CRLB, goodness of fit, SD of residual) | *CRLBs of GABA and Glu also provided in Table 2* |
| d. Sample Spectrum | Provided |
